# A novel cuproptosis-related long non-coding RNAs model that effectively predicts prognosis in hepatocellular carcinoma

**DOI:** 10.1101/2022.06.07.495148

**Authors:** Enmin Huang, Ning Ma, Tao Ma, Junyi Zhou, Weisheng Yang, Chuangxiong Liu, Zehui Hou, Shuang Chen, Zhen Zong, Bing Zeng, Yingru Li, Taicheng Zhou

## Abstract

**Background:** Cuproptosis has recently been considered a novel form of programmed cell death. To date, factors crucial to the regulation of this process remain unelucidated. Here, we aimed to identify long-chain non-coding RNAs (lncRNAs) associated with cuproptosis in order to predict the prognosis of patients with hepatocellular carcinoma (HCC).

**Methods:** Using RNA sequence data from The Cancer Genome Atlas Live Hepatocellular Carcinoma (TCGA-LIHC), a co-expression network of cuproptosis-related mRNAs and lncRNAs was constructed. For HCC prognosis, we developed a cuproptosis-related lncRNA signature (CupRLSig) using univariate Cox, lasso, and multivariate Cox regression analyses. Kaplan-Meier analysis was used to compare overall survival among high- and low-risk groups stratified by median CupRLSig score. Furthermore, comparisons of functional annotation, immune infiltration, somatic mutation, TMB (tumor mutation burden), and pharmacologic options were made between high- and low-risk groups.

**Results:** Our prognostic risk model was constructed using the cuproptosis-related PICSAR, FOXD2-AS1, and AP001065.1 lncRNAs. The CupRLSig high-risk group was associated with poor overall survival (hazard ratio = 1.162, 95% CI = 1.063– 1.270; p < 0.001). Model accuracy was further supported by receiver operating characteristic and principal component analysis as well as internal validation cohorts. A prognostic nomogram developed considering CupRLSig data and a number of clinical characteristics were found to exhibit adequate performance in survival risk stratification. Mutation analysis revealed that high-risk combinations with high TMB carried worse prognoses. Finally, differences in immune checkpoint expression and responses to chemotherapy as well as in targeted therapy among CupRLSig stratified high- and low-risk groups were explored.

**Conclusions:** The lncRNA signature constructed in this study is valuable in prognostic estimation in the setting of HCC.

## INTRODUCTION

With a 5-year survival rate of 18% and a median survival time of 1 year, liver cancer is the second most lethal tumor after pancreatic cancer (1). Hepatocellular carcinoma (HCC) accounts for about 80% of all primary liver tumors (2). Surgery, ablation, and orthotopic liver transplantation remain the most popular locoregional treatment options for HCC (3). However, as most HCC patients are diagnosed late in the illness and often suffer metastases on diagnosis, surgical resection is rarely a viable treatment option. Such patients can only be treated with systemic therapies, such as targeted therapy (4). Despite the availability of several tyrosine kinase inhibitors for first- and second-line treatment, overall survival (OS) in advanced HCC remains poor due to drug resistance and has not improved over the last decade (5). Although the recent FDA approval of immune checkpoint inhibitors (ICI) has transformed clinical management of HCC, only a small proportion of patients are sensitive to this therapy due to a lack of relevant selective biomarkers (6). As such, novel treatment modalities and prognostic markers warrant investigation to urgently improve patient outcomes.

Levels of copper, including the complex form of ceruloplasmin, are known to be significantly elevated in serum and tumors among cancer patients (7). Excess copper acts as a powerful oxidant, promoting the intracellular production of reactive oxygen species (ROS) and apoptosis (8). Malignant cells naturally possess higher basal ROS levels compared to normal cells (8) as they utilize mechanisms such as compensatory upregulation of NRF2 genes to counter increases in ROS resulting from copper accumulation (2). Thus, utilization of altered copper distribution to generate an intolerable increase of ROS stress in malignant cells warrants consideration as a potential anticancer strategy (7). Prior to the clinical utilization of spatial copper distribution for cancer treatment, however, copper metabolism genes and regulatory networks must first be known. For example, alterations in copper bioavailability have been investigated in preclinical studies of KRAS mutated tumors (9). Recently, researchers found that some cancer cells die when carrier molecules, such as FDX1, import substantial levels of copper into the cytoplasm (10). By blocking other alternative cell death pathways, this proved to be a specific kind of cell death, and further research revealed cells more reliant on mitochondria for energy production to be more sensitive to this copper-induced death, namely cuproptosis (10). Subsequent genome-wide CRISPR-Cas9 loss-of-function screens identified 10 genes involved in copper ionophore– induced death (10). The underlying regulatory roles and mechanisms of genes involved in cuproptosis in the setting of HCC, however, remain unclear.

Long non-coding RNAs (lncRNAs) are involved in a variety of biological processes. Several HCC-related lncRNAs were found to be abnormally expressed in tumor tissues and play important roles in shaping the tumor microenvironment via epigenetic regulation (11). Similarly, lncRNAs were reported to play crucial roles in the regulation of metabolism of metal ion homeostasis. Some 2564 lncRNAs were found to be significantly up-regulated, and 1052 down-regulated, in a recently constructed toxic milk mouse model of Wilson’s disease (WD), which is characterized by a mutated ATP7B gene that affects copper transport (12). The cytosolic lncRNA P53RRA was found to displace p53 from the G3BP1-p53 complex, resulting in increased intranuclear p53 retention and manifestation of ferroptosis, a similar ion-induced form of programmed cell death (13). Although the mechanism characterizing the lncRNA-mediated epigenetic regulation of ferroptosis has been widely investigated (14), the lncRNA regulatory network associated with cuproptosis remains almost completely unknown. Given that lncRNAs are involved in a wide range of biological processes including ferroptosis, their involvement in the regulation of cuproptosis is highly likely. Thus, identification of lncRNA transcriptional changes is critical in characterizing cuproptosis and its relevance in the setting of malignancy.

Here, we developed a cuproptosis-related lncRNA signature (CupRLSig) and demonstrated its adequacy in predicting HCC patient prognosis. Furthermore, we constructed a nomogram considering CupRLSig data as well as a number of clinical features and compared gene enrichment, mutations, immune cell infiltration, and potential responses to targeted therapy and immunotherapy among CupRLSig-defined high- and low-risk groups. This study highlights the cuproptosis regulatory network, the understanding of which is critical for improving the efficacy of individualized HCC treatment.

## MATERIALS AND METHODS

### Dataset and sample extraction

RNA-sequencing data (RNA-seq), clinical characteristics, and mutation data of HCC patients were obtained from The Cancer Genome Atlas - Live Hepatocellular Carcinoma Database (TCGA-LIHC, https://portal.gdc.cancer.gov/). Initially, data from 424 HCC patients were collected. Patients with incomplete follow-up data, survival < 30 days or lacking complete clinicopathological data were excluded from follow-up analysis; 343 patients were ultimately retained. The 19 cuproptosis-related genes, listed in Supplemental Table 1, were obtained from available literature (2, 9, 10, 15–17) reporting findings of gene manipulation studies either inducing or inhibiting cuproptosis.

### Identifying CupRLSig in predicting HCC patient prognosis

The absolute value of the Pearson correlation coefficient (> 0.4) and p < 0.05 were considered thresholds for the establishment of a cuproptosis-related mRNA-lncRNA co-expression network to identify lncRNAs relevant in cuproptosis. The network was visualized using a Sankey diagram generated by the R software package “ggalluvial.” The entire TCGA-LIHC sample was subsequently randomly divided into a training group and a validation group (Table 1); univariate Cox regression analysis was applied to determine whether these lncRNAs were associated with training group patient prognosis. A lasso regression analysis was additionally performed to avoid over-fitting and eliminate tightly correlated genes. Ten-fold cross-validation was employed to select the minimal penalty term (Lambda). These aforementioned lncRNAs were subsequently used to construct a multivariate Cox regression model and determine correlation coefficients. The model risk score formula obtained was as follows: risk score = explncRNA1×coef lncRNA1 + explncRNA2×coef lncRNA2 +…+ explncRNAi×coef lncRNAi. We termed this predictive lncRNA signature as CupRLSig. The risk score of each patient from the training, test and entire TCGA-LIHC groups was calculated, with HCC samples from all three groups divided into high- and low-risk groups based on training group median risk score value. Kaplan-Meier curves, risk curves, survival status, and heatmap analyses were employed to investigate whether the CupRLSig model effectively distinguishes patients of different risk levels. Model accuracy was quantified utilizing progression free survival (PFS), the concordance index (C-index), independent prognostic analysis, and the receiver operating characteristic (ROC) curve. The R software package “pheatmap” was used to visualize clinicopathological variables of high- and low-risk groups from the entire TCGA-LIHC sample set; the distribution of patients with varying risk scores was evaluated using principal component analysis (PCA) and visualized using the R software package “scatterplot3d.” Finally, stratified analysis was performed using various pathological parameters to determine whether the model’s distinction between high- and low-risk groups significantly correlated with other clinical parameters.

**Table 1.** Clinical characteristics of TCGA-LIHC sample training and test groups (n = 343).

| Covariates | Sub Type | Entire TCGA-LIHC (%) | Test Group (%) | Training Group (%) | p-value |
| --- | --- | --- | --- | --- | --- |
| Age | <=65 | 216(62.97%) | 103(60.23%) | 113(65.7%) | 0.3493 |
|  | >65 | 127(37.03%) | 68(39.77%) | 59(34.3%) |  |
| Gender | Female | 110(32.07%) | 59(34.5%) | 51(29.65%) | 0.3971 |
|  | Male | 233(67.93%) | 112(65.5%) | 121(70.35%) |  |
| Grade | G1 | 53(15.45%) | 27(15.79%) | 26(15.12%) | 0.3 |
|  | G2 | 161(46.94%) | 85(49.71%) | 76(44.19%) |  |
|  | G3 | 112(32.65%) | 54(31.58%) | 58(33.72%) |  |
|  | G4 | 12(3.5%) | 3(1.75%) | 9(5.23%) |  |
|  | Unknown | 5(1.46%) | 2(1.17%) | 3(1.74%) |  |
| Stage | Stage I | 161(46.94%) | 81(47.37%) | 80(46.51%) | 0.9079 |
|  | Stage II | 77(22.45%) | 39(22.81%) | 38(22.09%) |  |
|  | Stage III | 80(23.32%) | 38(22.22%) | 42(24.42%) |  |
|  | Stage IV | 3(0.87%) | 2(1.17%) | 1(0.58%) |  |
|  | Unknown | 22(6.41%) | 11(6.43%) | 11(6.4%) |  |
| T | T1 | 168(48.98%) | 86(50.29%) | 82(47.67%) | 0.5683 |
|  | T2 | 84(24.49%) | 42(24.56%) | 42(24.42%) |  |
|  | T3 | 75(21.87%) | 37(21.64%) | 38(22.09%) |  |
|  | T4 | 13(3.79%) | 4(2.34%) | 9(5.23%) |  |
|  | Unknown | 3(0.87%) | 2(1.17%) | 1(0.58%) |  |
| M | M0 | 245(71.43%) | 118(69.01%) | 127(73.84%) | 0.9551 |
|  | M1 | 3(0.87%) | 2(1.17%) | 1(0.58%) |  |
|  | Unknown | 95(27.7%) | 51(29.82%) | 44(25.58%) |  |
| N | N0 | 239(69.68%) | 111(64.91%) | 128(74.42%) | 0.9081 |
|  | N1 | 3(0.87%) | 2(1.17%) | 1(0.58%) |  |
|  | Unknown | 101(29.45%) | 58(33.92%) | 43(25%) |  |

The p-value is indicated for the one-way ANOVA test among the three groups.

### Construction of the nomogram

A nomogram was constructed using the R software packages “rms” and “regplot” for the prediction of HCC patient survival at 1-, 3-, and 5-years based on a combination of risk scores with other clinicopathological data. The calibration curve was used to evaluate whether predicted survival rate was consistent with actual survival rate. A patient was randomly selected to confirm the predictive utility of the nomogram.

### Functional enrichment analysis of differentially expressed genes and lncRNAs among high- and low-risk CupRLSig groups

Differentially expressed genes and lncRNAs among high- and low-risk CupRLSig groups were identified using the R software package “limma” with a log_2_ fold change absolute value greater than 1 and a false discovery rate (FDR) of <0.05. Functional enrichment analysis of the differentially expressed genes and lncRNAs was then performed using the Gene Ontology (GO) and the Kyoto Encyclopedia of Genes and Genomes (KEGG) databases.

### Analysis of somatic mutation data and tumor mutation burden (TMB)

The number of somatic non-synonymous point mutations in each sample was counted and visualized using the R software package “maftools” (18). The TMB was calculated as the number of somatic, coding, base replacement, and insert-deletion mutations discovered per megabase of genome using non-synonymous and code-shifting indels and a 5% detection limit. In addition, TMB was compared between high- and low-risk groups, and survival curves for TMB and risk score integration were plotted.

### Estimation of immune infiltration

The CIBERSORT algorithm (19) was used to estimate infiltration proportionality of 22 immune cell types in HCC samples. The Wilcoxon rank-sum test was used to determine whether there was a significant difference in immune cell proportions between low- and high-risk groups. Single-sample gene set enrichment analysis (ssGSEA) was performed using the R software package “GSVA” (20) to assess the activity of 13 immune-related functions and compare differences between the two groups.

### Potential relationship between CupRLSig and immunotherapy, chemotherapy, and target therapy

First, differential expression of 47 immune checkpoint genes in CupRLSig high- and low-risk groups was compared. The tumor immune dysfunction and exclusion (TIDE, http://tide.dfci.harvard.edu/) module was used to distinguish potential immunotherapy responses among groups. This module predicted anti-PD1 and anti-CTLA4 treatment responses based on patient pre-treatment genome transcriptional expression profiles. Further evaluation of the role of CupRLSig in predicting the therapeutic response of HCC involved calculation of the half-maximal inhibitory concentration (IC_50_) of commonly used chemotherapeutic as well as of targeted therapeutic drugs. The Wilcoxon signed-rank test and R software package “pRRophetic” were used to compare and visualize IC_50_ values in high- and low-risk groups.

### Statistical Analysis

The Kaplan-Meier method and log-rank test were used to compare OS and PFS among high- and low-risk group patients. The R software “survivalROC” package was used to construct ROC curves and calculate the area under the curve (AUC). The Kruskal–Wallis test was used to compare differences between groups and clinical data were analyzed using either chi-squared or the Fisher’s exact tests. Relationships between lncRNA expression, immune infiltration and immune checkpoint gene expression were assessed using Spearman or Pearson correlation coefficients. All statistical analyses were performed using R software (Version 4.1.2); a p-value < 0.05 was considered to indicate statistical significance.

## RESULTS

### Construction of the CupRLSig model

Figure 1 depicts the flow chart of the present study. First, Pearson correlation analysis identified 157 cuproptosis-related lncRNAs related to 14 cuproptosis genes considering a correlation coefficient > 0.4 and p < 0.05 (Figure 2A and Supplemental Table 2). The entire TCGA-LIHC sample was subsequently randomly divided into a training group and a validation group (Table 1). Univariate Cox regression analysis revealed a total of 27 lncRNAs to possess a prognostic correlation with the training group (Figure 2B). Following lasso regression analysis (Figure 2C and 2D), three lncRNAs were finally retained in the training group and used to construct a multivariate Cox regression model. The correlation between these three lncRNAs and 19 cuproptosis-related genes is shown in Figure 2E. We termed this lncRNA prediction signature as CupRLSig. The CupRLSig risk score formula was determined to be as follows: risk score = (0.2659×PICSAR expression) + (0.4374×FOXD2-AS1 expression) + (−0.3467×AP001065.1 expression). This formula was used to calculate the risk score for each patient and patients were divided into two risk groups based on training group median risk score. Finally, of the three training, test, and entire groups, 86, 80, and 166 patients, respectively, were assigned to the high-risk group; 86, 91, and 177 patients were assigned to the low-risk group (Figure 3A-3C). Kaplan-Meier analysis revealed a significantly shorter high-risk group OS as compared with the low-risk group among both datasets (Figure 3A-3C). Individual patient risk scores and survival statistics are detailed in Figure 3D-3I, with the number of deaths increasing as risk score increases. The expression status of three lncRNAs from each group is detailed in Figure 3J-3L.

**Figure 1.**
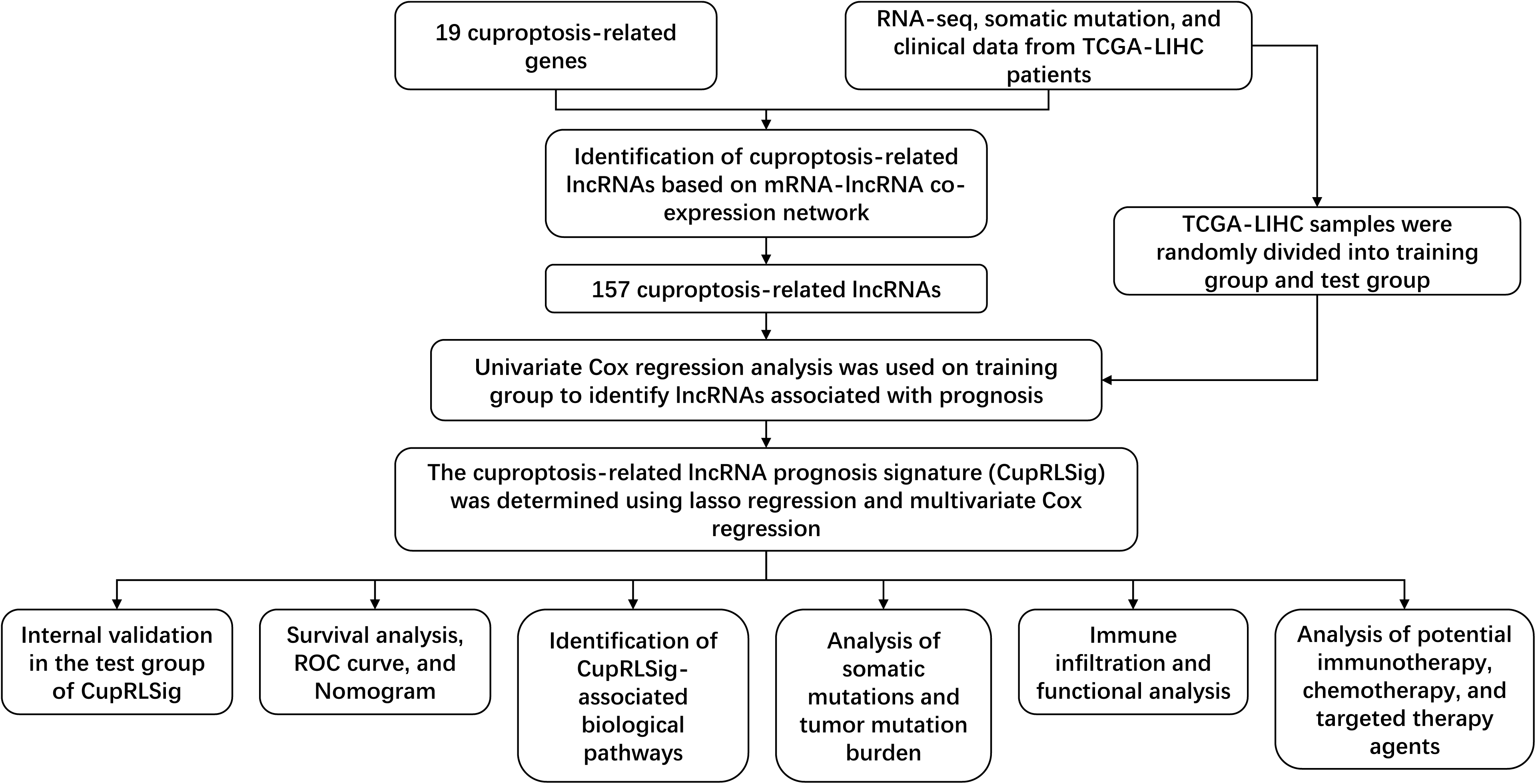
Study Flowchart. RNA-seq, RNA sequence; TCGA-LIHC, The Cancer Genome Atlas-Live Hepatocellular Carcinoma; lncRNAs, long non-coding RNAs; ROC, receiver operating characteristic.

**Figure 2.**
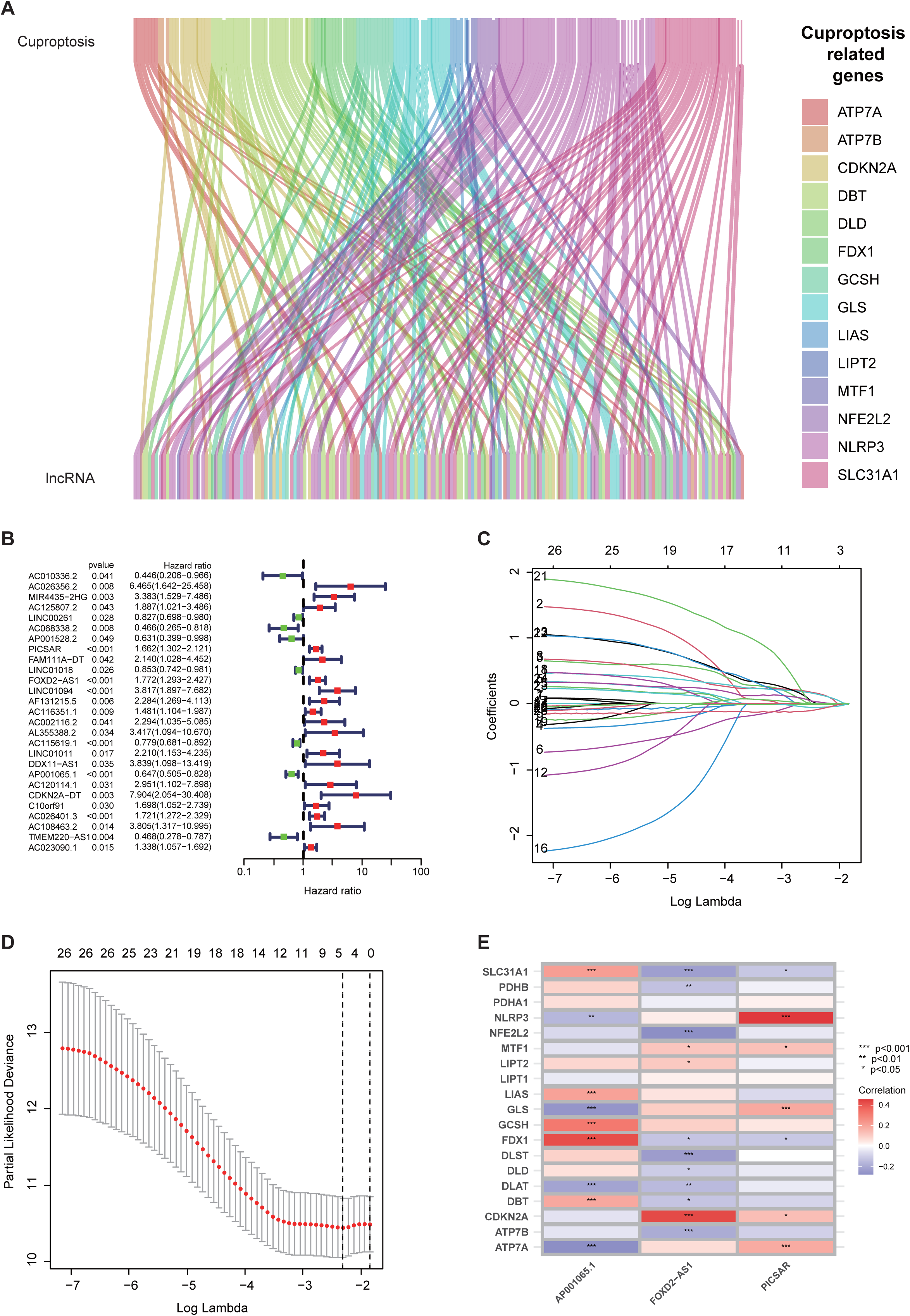
Construction of the CupRLSig model. (A) The Sankey diagram shows the associations between cuproptosis-related lncRNAs and mRNAs. (B) The Forest plot shows 27 lncRNAs with hazard ratios (95% confidence intervals) and p-values for their association with HCC prognosis based on univariate Cox proportional-hazards analysis. (C) Lasso coefficient profiles. (D) Selection of the tuning parameter (Lambda) in the lasso model by 10-fold cross-validation based on minimum criteria for overall survival. (E) A heatmap shows the correlation between the three lncRNAs incorporated into the CupRLSig model and 19 cuproptosis-related genes.

**Figure 3.**
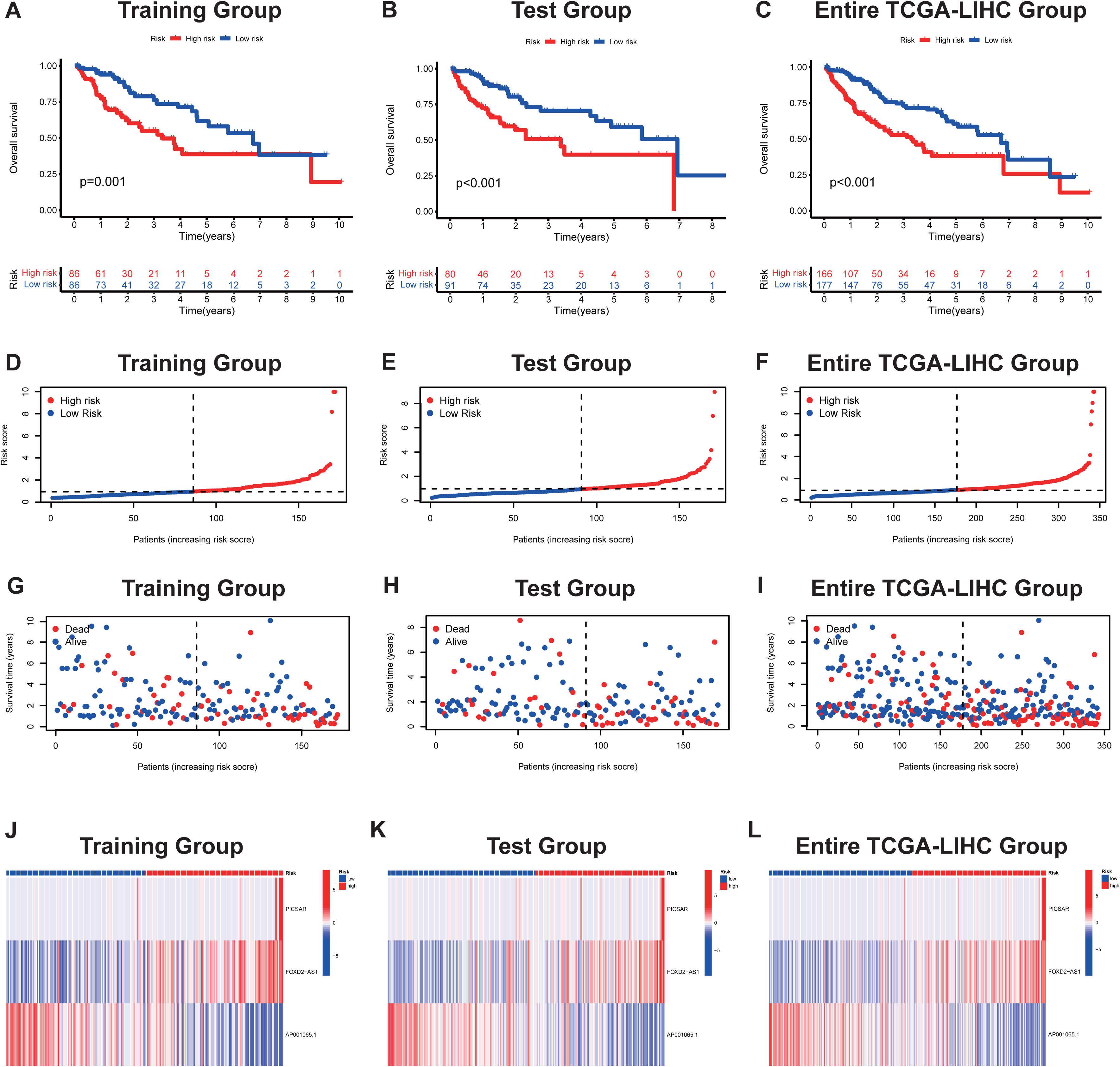
Internal validation for CupRLSig model overall survival determination for training, test, and entire TCGA-LIHC groups. Kaplan-Meier survival curves in the high- and low-risk groups stratified by median training group overall survival CupRLSig risk scores (A); test group data (B); and entire TCGA-LIHC group data (C). P-values were determined using the log-rank test. The risk curve is based on the risk score for each sample in training (D), test (E) and entire TCGA-LIHC (F) groups, where red and blue dots indicate high- and low-risk samples, respectively. The scatter plot is based on the survival status of each sample from training (G), test (H) and entire TCGA-LIHC (I) groups, where red and blue dots indicate death and survival, respectively. (J-L) Heatmaps detail expression levels of the three CupRLSig lncRNAs in each group. TCGA-LIHC, The Cancer Genome Atlas-Live Hepatocellular Carcinoma.

### Evaluate the accuracy of the CupRLSig model

We further evaluated the PFS of 343 HCC patients using data downloaded from http://xena.ucsc.edu/ to assess prediction accuracy of our CupRLSig prognostic model among HCC patients. High-risk patients were noted to have significantly shorter PFS (p = 0.001; Figure 4A). The C-index revealed the model’s prognostic prediction performance to be comparable to disease stage (Figure 4B). Univariate and multivariate Cox regression analyses revealed CupRLSig risk score to be an independent prognostic factor (Figure 4C and 4D); its AUC of 0.741 was found to be a better predictor of HCC prognosis as compared to other clinicopathological variables (Figure 4E). 1-, 3-, and 5-year ROC AUCs were 0.741, 0.636, and 0.649, respectively, indicating that CupRLSig exhibited good prognostic performance (Figure 4F).

**Figure 4.**
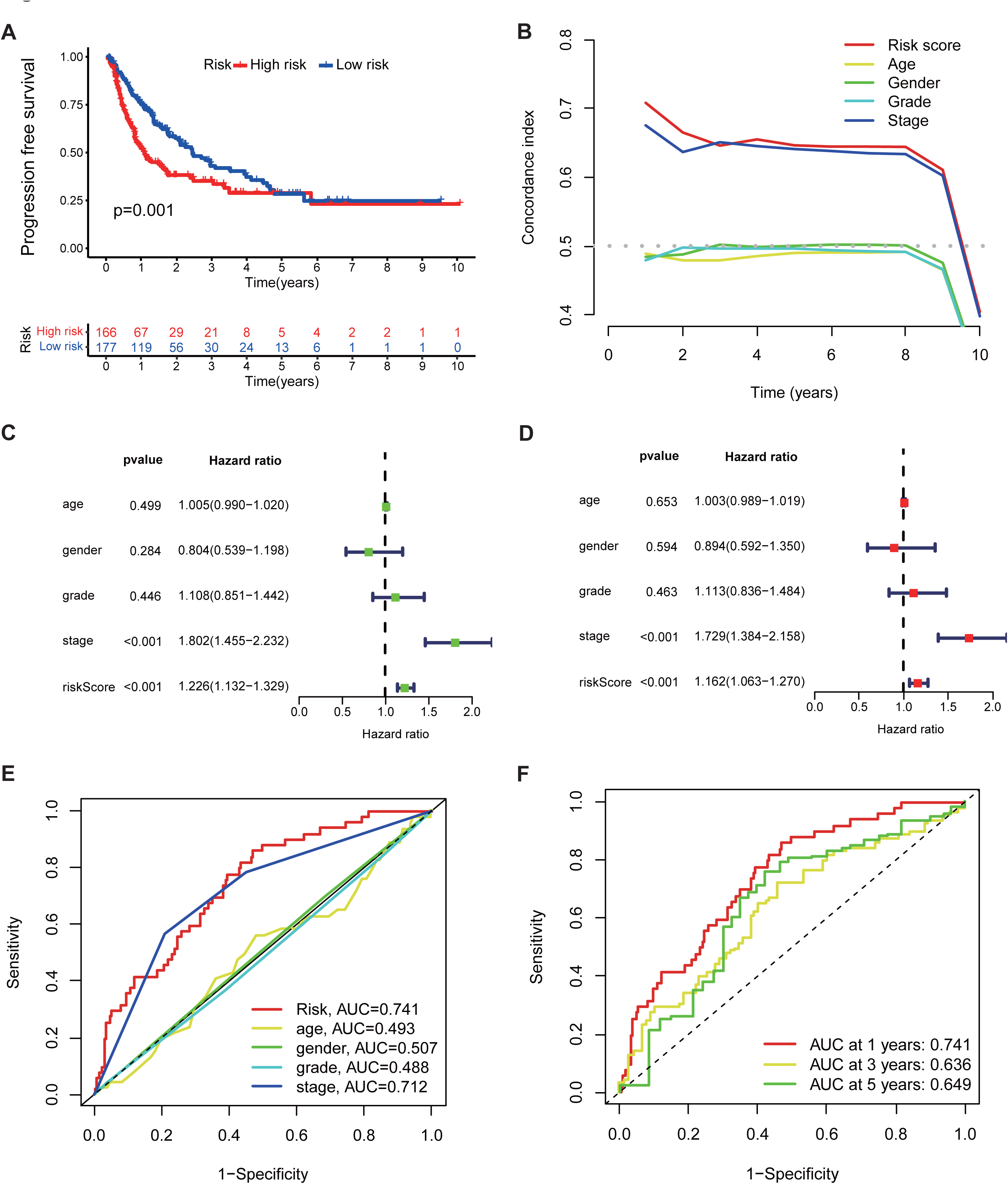
Evaluation of CupRLSig model predictive accuracy using the entire TCGA-LIHC group. (A) Kaplan–Meier curves for progression-free survival in high- and low-risk groups stratified by median of CupRLSig risk scores. (B) The concordance index curves depict CupRLSig risk scores and other clinical parameters relevant to predicting HCC patient prognosis. Forest plots for univariate (C) and multivariate (D) Cox proportional-hazard analysis for determination of the independent prognostic value of the CupRLSig risk score. (E) ROC curve of the CupRLSig risk score and other clinicopathological variables. (G) Time-dependent ROC curves for 1-, 3-, and 5-year survival for the CupRLSig signature. TCGA-LIHC, The Cancer Genome Atlas-Live Hepatocellular Carcinoma. ROC, receiver operating characteristic. AUC, area under the curve.

Expression levels of the three lncRNAs from the CupRLSig model, as well as clinicopathological factors, are detailed in Figure 5A. The PCA of whole genes, cuproptosis genes, cuproptosis lncRNAs and risk lncRNAs from the CupRLSig model was performed to distinguish between high- and low-risk patients (Figure 5B-5E). The CupRLSig (Figure 5E) model was found to effectively distinguish among low- and high-risk groups, underscoring the accuracy of the model.

**Figure 5.**
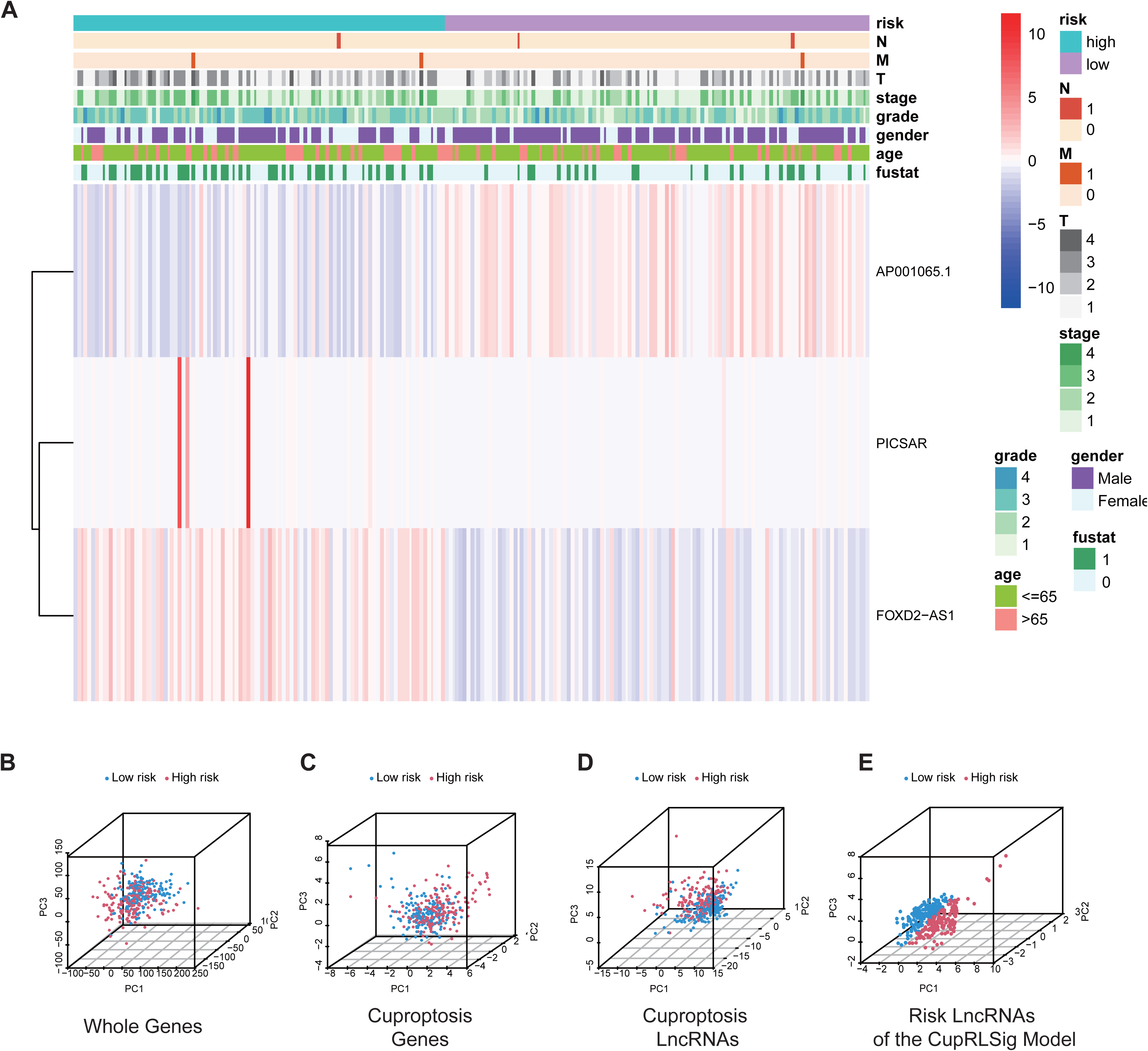
Visualization of expression levels of the three CupRLSig model component lncRNAs based on clinicopathological variable stratification and principal component analysis (PCA) of different gene sets performed for classification of patient risk. (A) A heatmap of the three lncRNAs and clinicopathological variables was constructed for high- and low-risk groups. PCA of low- and high-risk groups based on (B) whole-genome genes, (C) cuproptosis-related genes, (D) cuproptosis-related lncRNAs, and (E) CupRLSig model risk lncRNAs. Patients with high risk scores are denoted by red, while those with low risk scores are denoted by blue. N, lymph node metastasis; M, distant metastasis; T, tumor.

Whether CupRLSig had prognostic value in subgroups with different clinicopathological parameters was also assessed (Figure 6A to 6J). Significant correlations between risk score and age (Figure 6A and 6B), sex (Figure 6C and 6D), tumor grade (Figure 6E and 6F), tumor stage (Figure 6G and 6H), and T stage (Figure 6I and 6J) were noted when assessing correlations among risk score and clinicopathological factors. The number of M and N stage subgroup cases was too small for evaluation. As such, the CupRLSig risk score was found to be an independent prognostic risk factor for HCC patients.

**Figure 6.**
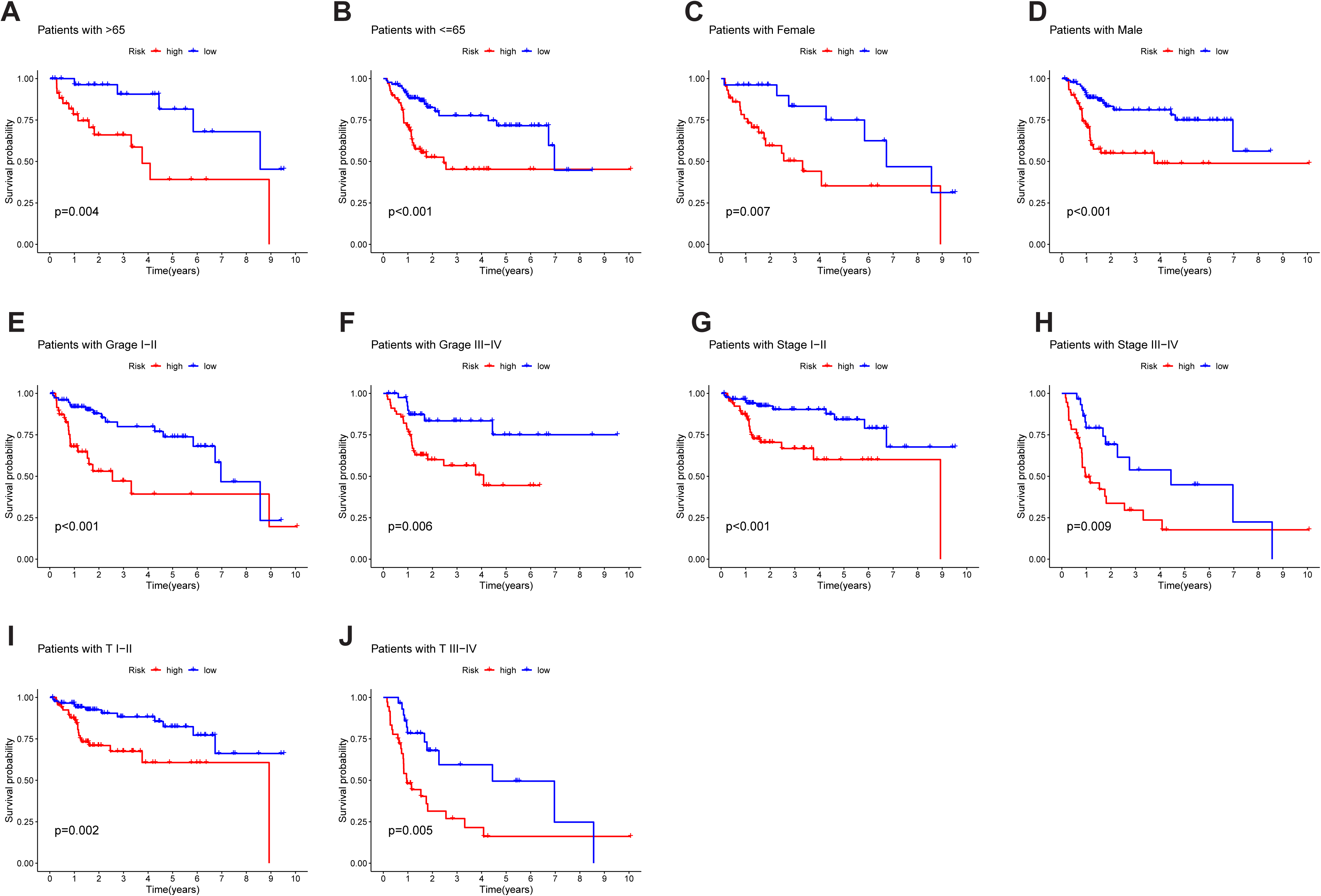
Kaplan-Meier survival curves for high- and low-risk patient groups sorted by clinicopathological variables. (A-B) Age; (C-D) Sex; (E-F) Grade; (G-H) Overall stage; (I-J) T stage. T, tumor.

### Construction of a predictive nomogram

The CupRLSig risk score, in combination with other clinicopathological factors, was used to develop a nomogram to guide clinical assessment of prognosis and estimate HCC patient 1-, 3-, and 5-year survival probability (Figure 7A). The 53rd patient was chosen for randomly evaluating the predictive utility of the nomogram. As shown in Figure 7A, the corresponding score of the 53rd patient was 175 points; the 5-year survival rate was 0.642, the 3-year survival rate was 0.738, and the 1-year survival rate was 0.875. The nomogram was found to accurately estimate mortality rate (Figures 7B).

**Figure 7.**
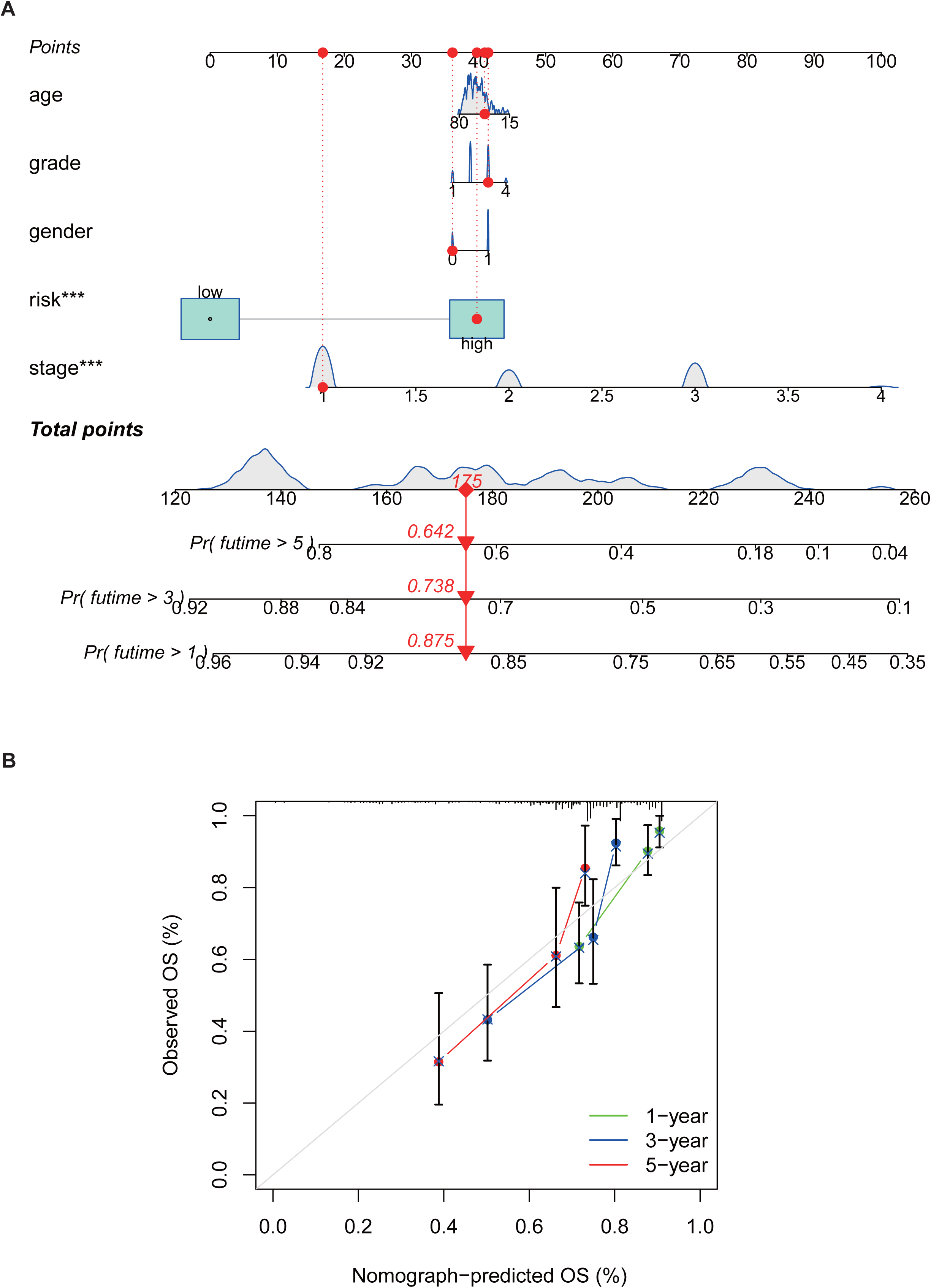
Nomogram construction and verification. (A) A nomogram combining clinicopathological parameters and risk scores predicts 1-, 3-, and 5-year survival probabilities of HCC patients. The multivariate Cox proportional hazard analysis was used to determine each parameter’s independent prognostic value. The red dots, diamonds, triangles, and dashed lines represent the 53rd patient randomly selected for the nomogram illustration. Calibration curves assess the consistency between observed actual and nomogram-predicted overall survival at (B) 1-, (C) 3-, and (D) 5-years. OS, overall survival.

### Identification of biological pathways linked to CupRLSig

The R software “enrichplot” package was used for gene set functional annotation of differentially expressed genes and lncRNAs (n = 523, Supplemental Table 3) among high- and low-risk HCC groups. The five biological processes found considering GO to possess the highest enrichment were mitotic nuclear division, mitotic sister chromatid segregation, nuclear division, chromosome segregation, and sister chromatid segregation (Figure 8A). The five cellular components found to possess the highest enrichment were condensed chromosomes, kinetochores, spindle, chromosomes, and condensed chromosomes (Figure 8A). Finally, the most enriched molecular functions were found to be steroid hydroxylase activity, oxidoreductase activity, microtubule binding, aromatase activity, and tubulin binding (Figure 8A). The five most enriched KEGG pathways were found to be retinol metabolism, cytochrome P450 drug metabolism, cytochrome P450 xenobiotic metabolism, the cell cycle, and chemical carcinogenesis-DNA adducts (Figure 8B).

**Figure 8.**
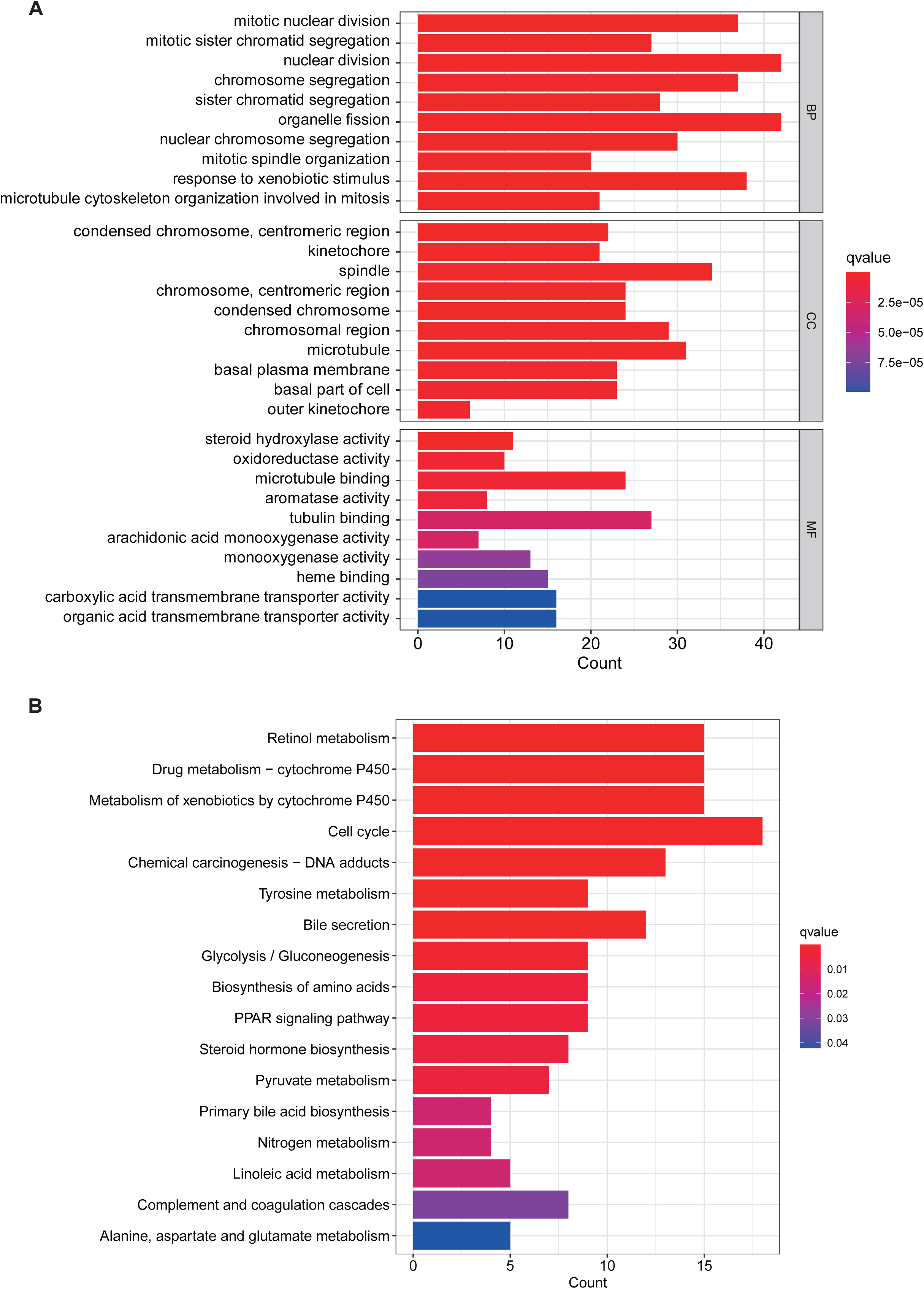
Gene set functional annotation of differentially expressed genes and lncRNAs in high- and low-risk HCC groups. (A) In biological process GO terms, differentially expressed genes and lncRNAs were found to be most enriched in mitotic nuclear division, mitotic sister chromatid segregation, nuclear division, chromosome segregation, and sister chromatid segregation; in the five cellular components of condensed chromosomes, kinetochores, spindles, chromosomes, and condensed chromosomes; and in the five molecular functions of steroid hydroxylase activity, oxidoreductase activity, microtubule binding, aromatase activity, and tubulin binding. (B) Differentially expressed genes and lncRNAs were found to be most enriched in the five KEGG pathways of retinol metabolism, cytochrome P450 drug metabolism, cytochrome P450 xenobiotic metabolism, cell cycle, and chemical carcinogenesis-DNA adducts. GO, gene ontology; KEGG, Kyoto encyclopedia of genes and genomes; BP, biological process; CC, cellular component; MF, molecular function.

### The relationship between CupRLSig risk scores and somatic mutation and TMB

Somatic mutations in low- and high-risk subgroup patients were assessed separately (Figure 9A and 9B); TP53 (36% vs. 17%) had a higher rate of somatic mutation in the high-risk group, while CTNNB1 (30% vs. 20%) and TTN (25% vs.20%) had a higher rate of somatic mutation in the low-risk group. Furthermore, although no difference in TMB between the two groups (Figure 9C) was found, survival time of patients with higher TMB was significantly reduced (Figure 9D). High TMB among high-risk group patients led to an even worse prognosis (Figure 9E), highlighting a significant synergistic effect between these two indicators.

**Figure 9.**
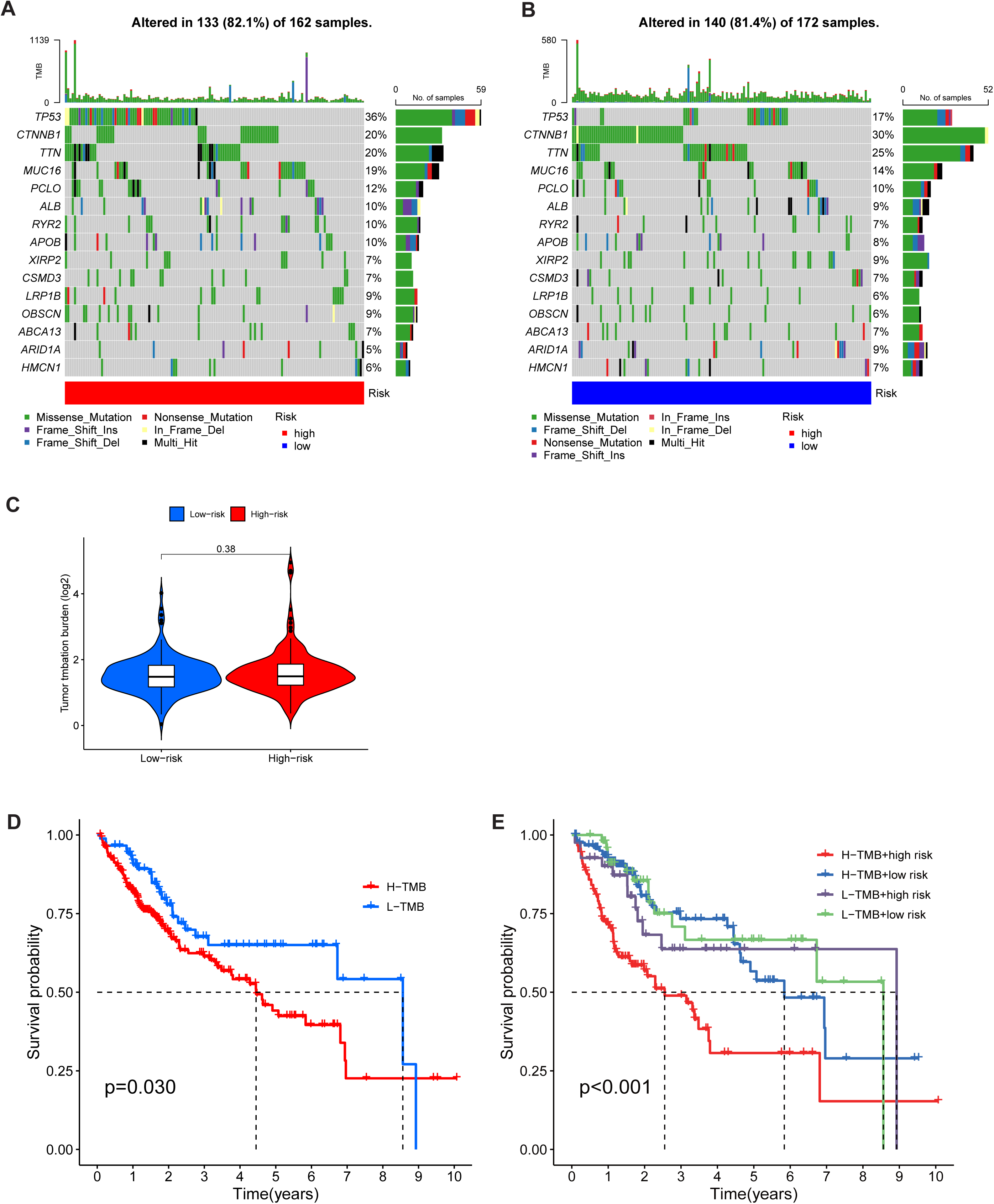
The relationship between CupRLSig risk scores and somatic mutation and tumor mutation burden (TMB). The waterfall plots showing somatic mutations of the most significant 15 genes among high-risk (A) and low-risk (B) HCC patients. (C) TMB comparison between low- and high-risk subgroups. (D) Kaplan-Meier curves for high- and low-TMB groups. (E) Subgroup analyses for Kaplan-Meier curves of patients stratified by TMB and risk scores. The p-value is representative of the ANOVA test among subgroups.

### Immune infiltration in different risk subgroups

The CIBERSORT algorithm revealed that the infiltration ratio of M2 macrophages (p = 0.007), resting mast cells (p = 0.002), monocytes (p = 0.002), and activated NK cells (p = 0.032) in the low-risk group was significantly greater as compared to the high-risk group (Figure 10A). Ratios of resting NK cells (p = 0.018), regulatory T cells (Tregs; p = 0.021), CD4 memory activated T cells (p = 0.025), and M0 macrophages (p = 0.007) exhibited the opposite pattern (Figure 10A). Scores of immune functions such as the C-C chemokine receptor (CCR), check points, and major histocompatibility complex (MHC) class I were significantly higher in high-risk group patients as compared to those in the low-risk group, although response to interferon type II exhibited an opposite pattern (Figure 10B). These findings revealed differences in immune infiltration among the two groups. As immunotherapy is understood to depend on the pre-existence of a “hot” immune microenvironment (21), such differences highlight the potential of immunotherapy.

**Figure 10.**
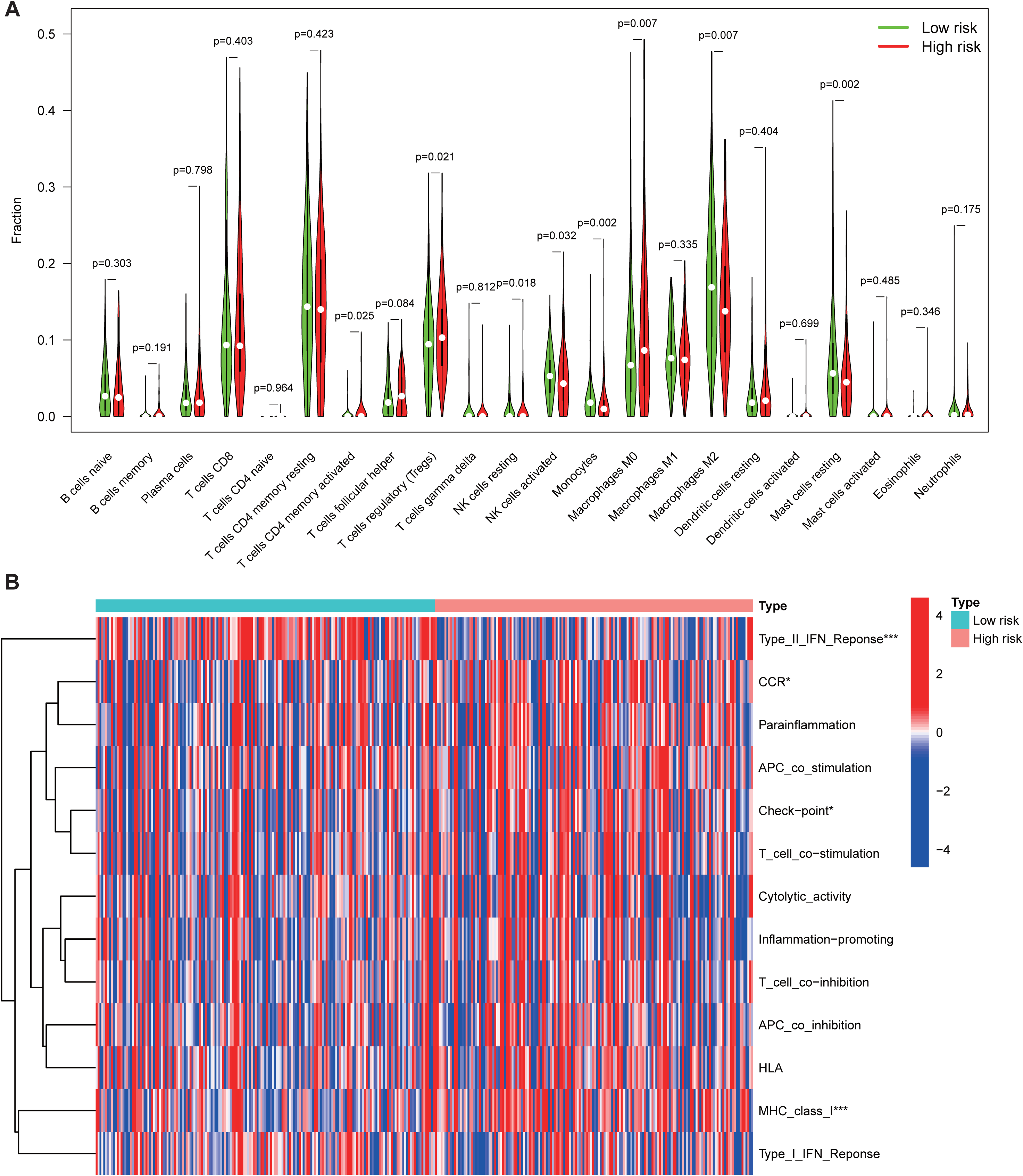
Immune cell infiltration and immune-related functions in different risk groups. (A) The violin plot shows whether there were significant differences in immune infiltration among 22 types of cells between high- and low-risk subgroups. (B) The heatmap shows whether there were significant differences in 13 immune-related functions between high- and low-risk subgroups. NK, natural killer; CCR, C-C chemokine receptor; APC, antigen-presenting cell; HLA, human leukocyte antigen; MHC, major histocompatibility complex; IFN, interferon. *p < 0.05; ***p < 0.001.

### Potential relationship between CupRLSig and immunotherapy, chemotherapy, and targeted therapy in HCC

Some relevant 28 genes were found to differ in expression levels between high- and low-risk groups out of a total of 47 immune checkpoints evaluated (Figure 11A). Immunotherapy markers such as CD276, CTLA-4, and PDCD-1, currently widely in clinical use, were found to be markedly elevated in the high-risk group (Figure 11A), implying potential immunotherapeutic responses in high-risk patients. Moreover, when the online software “TIDE” was used to predict the outcome of cancer patients treated with anti-PD1 or anti-CTLA4, a higher TIDE score was found in the low-risk group as compared to the high-risk group (Figure 11B). Importantly, a higher TIDE score suggests a greater likelihood of tumor immune escape and a poorer response to immunotherapy. Considering immune infiltration, checkpoint gene expression and the TIDE score, cuproptosis-related high-risk HCC patients are likely to respond better to immunotherapy.

**Figure 11.**
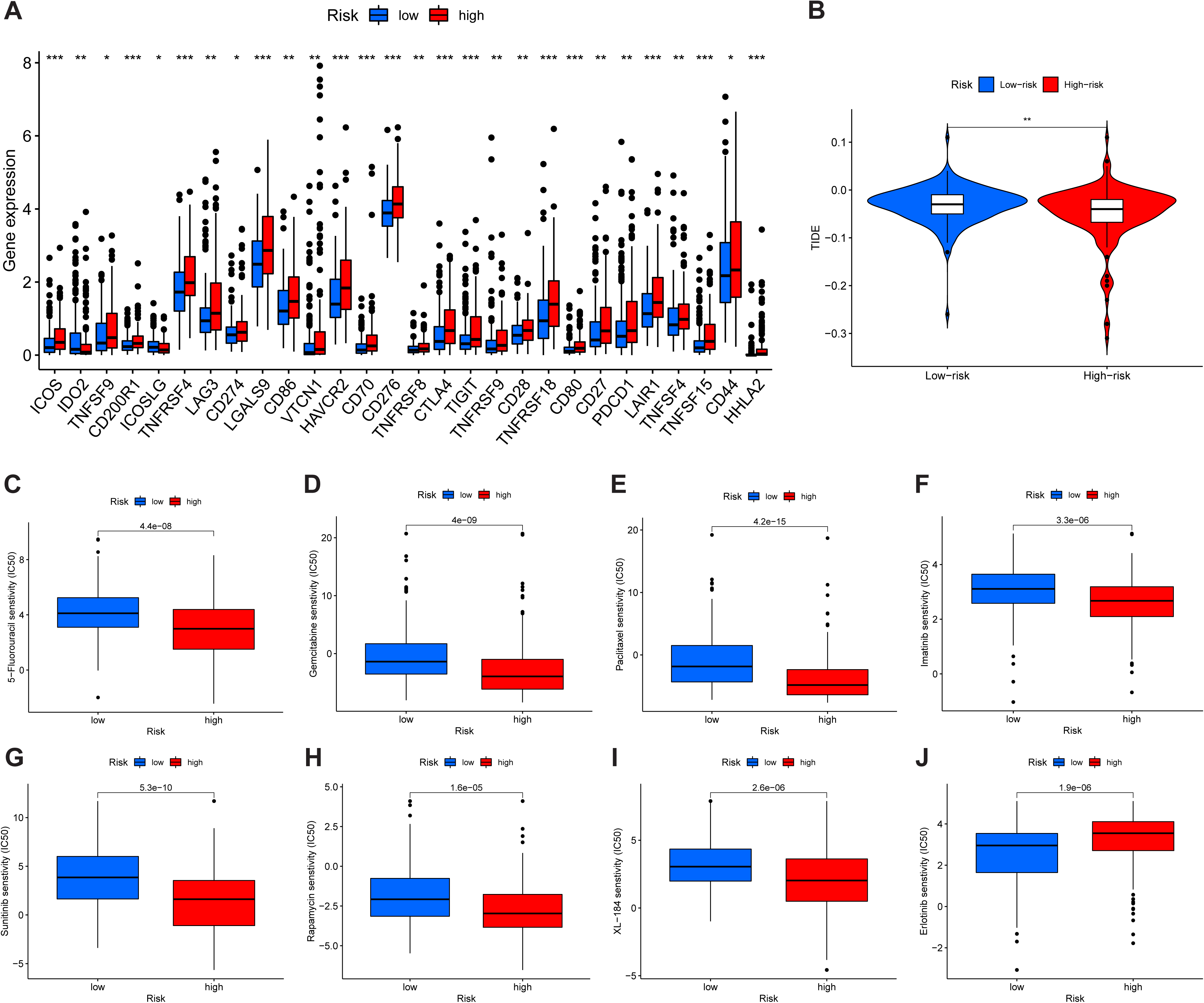
Comparison of Immune checkpoints, tumor immune dysfunction, and exclusion module (TIDE) scores, and chemotherapy and targeted therapy drug efficacy in high- and low-risk groups. (A) Expression of 28 immune checkpoint genes differs between the high- and low-risk groups. Red and blue boxes represent high- and low-risk patients, respectively. (B) Online software TIDE predicts HCC subgroup outcomes treated with either anti-PD1 or anti-CTLA4. A higher TIDE score suggests a greater likelihood of tumor immune escape and a poorer response to immunotherapy. The IC50 values for (C) 5-Fluorouracil, (D) Gemcitabine, (E) Paclitaxel, (F) Imatinib, (G) Sunitinib, (H) Rapamycin, (I) XL-184 (Cabozantinib), and (J) Erlotinib in high- and low-risk groups. IC50, half-maximal inhibitory concentration. *p < 0.05; **p < 0.01; ***p < 0.001; ns, non-significant.

Finally, the relationship between CupRLSig risk score and efficacies of chemotherapy and targeted therapy for HCC were evaluated. Most drugs commonly used in preclinical and clinical systemic therapy for HCC, such as 5-fluorouracil (Figure 11C), gemcitabine (Figure 11D), paclitaxel (Figure 11E), imatinib (Figure 11F), sunitinib (Figure 11G), rapamycin (Figure 11H), and XL-184 (cabozantinib, Figure 12I) were found to be more efficacious in the low-risk group; erlotinib (Figure 12J), an exception, was more efficacious in the high-risk group. Taken together, our findings underscore the potential that CupRLSig possesses in the future clinical development of personalized treatment strategies.

## DISCUSSION

Widespread hepatitis B vaccination in China has led to a gradual decline in HCC incidence, from 29.2/100,000 in 1998 to 21.9/100,000 in 2012 (22). However, HCC prognosis remains poor, in large part due to a lack of therapeutic and prognostic biomarkers. Markers currently considered in clinical practice, such as AFP, can be used as diagnostic markers or for monitoring recurrence, but they do not provide treatment or prognostic data (23). The combination of several biomarkers into a single model improves both therapeutic and prognostic prediction accuracy as compared to a single biomarker (24).

Serum and tissue copper levels are known to be elevated in the setting of various malignancies, with such elevation being directly related to cancer progression (7). As such, we hypothesized that abnormal expression of genes relevant to the copper metabolism pathway can serve as prognostic and therapeutic markers in the setting of HCC. Cuproptosis, a form of programmed cell death recently identified to result from the binding of accumulated intracellular copper to aliphatic components of the tricarboxylic acid cycle, causes lipoacylated protein aggregation and loss of iron-sulfur cluster proteins (10). Although many genes pivotal in cuproptosis have been identified, the overall regulatory landscape of this process in HCC remains unclear. Here, we incorporated signatures of three cuproptosis-related lncRNAs to develop a CupRLSig signature capable of addressing both cuproptosis and HCC prognosis.

Based on the ROC curve, CupRLSig was found to exhibit adequate predictive utility in the evaluation of OS among HCC patients. In addition, our novel nomogram improves clinical decision-making and has the potential to guide development of treatment strategies. In the CupRLSig model, both FOXD2-AS1 and PICSAR were previously identified as oncogenes in HCC, where FOXD2-AS1 aggravates HCC tumorigenesis by regulating the miR-206/MAP3K1 axis (25) while PICSAR accelerates disease progression by regulating the miR-588/PI3K/AKT/mTOR axis (26). However, there is a lack of research investigating the prognostic value of AP001065.1 and the magnitude of its involvement in cuproptosis and further study of this lncRNA is warranted.

This study also explored the important relationship between cuproptosis and treatment decisions for managing HCC. Endogenous oxidative stress levels are known to be elevated in a variety of tumors, likely due to a combination of active metabolism, mitochondrial mutations, cytokine activity, and inflammation (7). Under constant oxidative stress, cancer cells tend to make extensive use of adaptive mechanisms and may deplete intracellular ROS buffer capacity (7). Thus, increased copper levels in cancer cells, as well as the resulting increase in oxidative stress, present a novel cancer-specific therapeutic strategy. The liver is the most important organ for copper metabolism, with the biliary tract excreting 80% of copper ions (27). The induction of cuproptosis in the setting of HCC thus offers a basis for effective management of this illness. Application of such a concept to preclinical studies first requires a detailed understanding of cuproptosis pathway regulatory gene expression in HCC patients. Investigation of a WD mouse model revealed that ATP7B-deficient hepatocytes, such as those found in WD patients, activate autophagy in response to copper overload to prevent copper-induced apoptosis (15). Inhibition of the autophagy pathway and consequent further copper overload and elevated ROS thus likely activates the cuproptosis pathway and leads to the death of such copper-rich tumor cells. Interestingly, efficacy of chemotherapeutic agents designed to induce ROS, such as paclitaxel, differs between patients in high- and low-risk groups as defined by the CupRLSig model. The CupRLSig model was additionally shown to have a relationship with the HCC immune microenvironment. According to CupRLSig stratification, expression of most immune checkpoints, activation of immune pathways and infiltration of immune cells were greater in the high-risk group as compared to the low-risk group, while TIDE score was noted to exhibit an opposite pattern. These findings suggest that high-risk patients have more to benefit from immunotherapy. Taken together, this study confirms CupRLSig to possess utility as an adjunctive selection tool for pharmacotherapy.

There were several limitations to this study. First, only TCGA data sets were utilized. Use of additional external data, such as from the Gene Expression Omnibus (GEO), should be considered in future studies to further confirm predictive utilities of CupRLSig. Second, owing to a lack of complete data, prognostic factors such as surgical data were not considered for nomogram construction. This may have affected the accuracy of the model. Third, functional studies are required to better understand molecular mechanisms associated with effects of cuproptosis-related lncRNAs.

In conclusion, this study describes a novel CupRLSig lncRNA signature, also included in our nomogram, useful in predicting HCC prognosis. Importantly, CupRLSig likely also predicts the level of immune infiltration and potential efficacy of tumor immunotherapy, chemotherapy, and targeted therapy.

## DATA AVAILABILITY STATEMENT

Publicly available datasets were analyzed in this study. These data can be found here: https://portal.gdc.cancer.gov/repository.

## ETHICS STATEMENT

Not applicable.

## AUTHOR CONTRIBUTIONS

TZ, YL, BZ, ZZ, and SC conceived the study and its design, and provided administrative support. EH, NM, and TM were involved in data analyses and wrote, reviewed, and edited the manuscript. JZ, WY, CL, and ZH contributed data analysis and reviewed the manuscript. All authors read and approved the final manuscript. All authors contributed to the article and approved the submitted version for publication.

## ACKNOWLEDGMENTS

The authors would like to thank The Cancer Genome Atlas (TCGA) for providing useful RNA-seq data with detailed accompanying clinical information (https://tcga-data.nci.nih.gov/tcga/). The authors would like to thank Charlesworth Author Services and Guangzhou Vengene Technology Co., Ltd for their assistance with language editing.

---

**Supplemental Table 1.** Cuproptosis-related genes.

**Supplemental Table 2.** Cuproptosis mRNA and lncRNA network.

**Supplemental Table 3.** Differentially expressed genes and lncRNAs (n=523) among high- and low-risk HCC groups.

